# Insect-specific Alphamesonivirus-1 (*Mesoniviridae*) in lymph node and lung tissues from two horses with acute respiratory syndrome

**DOI:** 10.1101/2024.11.10.622896

**Authors:** Lucija Jurisic, Heidi Auerswald, Maurilia Marcacci, Francesca Di Giallonardo, Laureen M. Coetzee, Valentina Curini, Daniela Averaimo, Ayda Susana Ortiz-Baez, Cesare Cammà, Giovanni Di Teodoro, Juergen A. Richt, Edward C. Holmes, Alessio Lorusso

## Abstract

As members of the RNA virus order *Nidovirales* include those that infect hosts ranging from marine invertebrates to terrestrial mammals, understanding their emergence, host range and disease potential is of clear importance. The *Mesoniviridae* are a recently documented family of viruses within the *Nidovirales*. To date, mesoniviruses have only been associated with the infection of arthropods, particularly mosquitoes. Herein, we report the first detection of a mesonivirus – Alphamesonivirus-1 – in mammals. Specifically, we utilized genomic and histological techniques to identify the presence of Alphamesonivirus-1 in lung and lymph node tissues of two horses that succumbed to an acute respiratory syndrome. Notably, no other pathogens typically associated with respiratory disease in horses were detected in these samples. Counter to the previous contention that mesoniviruses only infect insects, our findings suggest a potentially broader host range and cross-species transmission of these viruses. The genome sequences of Alphamesonivirus-1 obtained from the two horses were closely related to those from a local *Culex* mosquito pool as well as an Alphamesonivirus-1 previously in identified Italy, suggestive of ongoing local transmission. The discovery of Alphamesonivirus-1 in tissues from diseased horses not only challenges current understandings of mesonivirus host range, but prompts further investigation into the role of insect-specific viruses in mammalian disease processes. Our results emphasize the importance of considering atypical pathogens in cases of unexplained animal deaths and suggest a potential zoonotic threat posed by previously overlooked viral families.

**IMPORTANCE:** Alphamesoniviruses, members of the *Mesoniviridae* family, have long been considered insect-specific viruses with no known association with vertebrate hosts. Herein, we describe the first detection of Alphamesonivirus-1 in mammals, marking a significant expansion of the known host range for this newly described virus family. Using detailed molecular and histological analyses we identified Alphamesonivirus-1 in lung and lymph node tissues of two horses that presented with an acute respiratory syndrome. Our findings indicate that Alphamesoniviruses may possess a broader host range than previously believed and could potentially induce severe disease in mammals. This unexpected host jump not only challenges existing knowledge on the ecology of mesoniviruses, but suggests that insect-specific viruses may pose a previously unrecognized health risk to vertebrates, including domesticated animals. These insights prompt the need for increased surveillance of atypical pathogens, especially in cases of unexplained respiratory illness, and may have implications for zoonotic disease emergence.

## INTRODUCTION

The emergence and global spread of SARS-CoV-2 has highlighted the capacity of viruses from the RNA virus order *Nidovirales* (i.e., coronaviruses) to jump species boundaries, occasionally resulting in outbreaks of infectious disease^1^. In most cases, such virus host jumps occur between closely related host taxa, such as different mammalian species, in part reflecting conserved virus-cell receptor relationships. SARS-CoV-2 has also been characterized by multi-species host jumping, with the virus passing from humans to a diverse array of mammalian species^2^. As a consequence, understanding the exact nature of host-pathogen interactions, including the barriers to successful cross-species transmission, is of considerable research interest^3^. Fortunately, our knowledge of the host range of many viruses has been greatly enhanced by the increasing use of metagenomic sequencing, which enables the entire viromes of species to rapidly documented. In addition, metagenomic sequencing has advanced diagnostics, enabling the identification and characterization of poorly described animal pathogens. However, despite our expanded knowledge of the virosphere, there has been a general neglect of the *Nidovirales* aside from the mammalian coronaviruses, including their prevalence, host range and frequency of host jumps, and their capacity to cause disease.

Respiratory problems are common in horses and are often diagnosed as a cause of poor athletic performance. However, the basic diagnostic techniques of the equine respiratory tract examination are not always sufficient for a complete diagnosis of the disease, its exacerbation, remission, or response to treatment. Of the different causes that might lead to respiratory system problems, infections are the most common disorders. This is particularly so with racehorses, in which respiratory system infections are often cited as the second most common reason for horses failing to train (Ainsworth and Hackett, 2004). Pathogens of the greatest concern in horses are influenza A viruses (AIV), Equine herpesvirus 1 and 4 (EHV1, EHV4), *Streptococcus zooepidemicus, Streptococcus pneumoniae, Streptococcus equi subsp. equi, Rhodococcus equi*, and *Pasteurella spp*.^4–6^. In addition, horses are susceptible to a plethora of viruses transmitted by biting arthropods, including mosquitoes, midgets, flies, ticks, as well as parasites^7^.

The *Mesoniviridae* are a newly assigned family of viruses within the *Nidovirales*. Unlike other nidoviruses, mesoniviruses are considered insect-specific viruses (ISVs) mainly found in mosquitoes and are not known to infect vertebrate cells. This family contains only one subfamily, *Hexponivirinae*, that currently contains a single genus – *Alphamesonivirus* – and nine subgenera (https://ictv.global/taxonomy).

The genome of Alphamesonivirus-1 (*Alphamesonivirus cavallyense*, subgenus *Namcalivirus*) contains seven open reading frames (ORFs), with ORF1a and -1b located at the 5′ end, encompassing two-thirds of the genome, and the smaller ORF2a, -2b, -3a, -3b, and -4 occupying the 3′-proximal end of the genome. The five major 3′-ORFs are predicted to encode a spike (S) glycoprotein (in ORF2a), a nucleocapsid (N) protein (in ORF2b), two proteins with membrane-spanning regions (in ORF3a and -3b), and a small protein with unknown function (in ORF4)^8,9^.

Alphamesonivirus-1 still comprises most of the mesoniviruses identified and was initially considered the prototype species of the family (https://ictv.global/taxonomy). The family was first represented by two closely related viruses: Cavally virus (CavV), isolated in Ivory Coast and initially named as a first insect-associated nidovirus^8^, and Nam Dinh virus (NDiV) isolated in Vietnam^10^, with the new family *Mesoniviridae* proposed the following year^11^. Mesoniviruses have only been identified from naturally infected mosquitoes^12,13^ and hence are considered to be ISVs^14,15^ in a similar manner to insect-specific flaviviruses^16^ and mosquito-associated bunyaviruses^17^. Indeed, to date, mesoniviruses have only been associated with invertebrates, with no reports in vertebrates^16,18^. Of note, a growing number of mesonivirus species have been identified from mosquitos collected in the Americas^19^, Asia^20^, Africa^12^, and Australia^21^, suggesting a near global distribution. Additionally, a mesonivirus was identified from *Aphis citricidus* aphids collected in China in 2012^22^, while a mesoni-like virus has been detected in an obligate fungal pathogen – *Leveillula Taurica* – in Italy^23^. Hence, the host range of mesoniviruses is likely to be far broader than currently known.

Here, using a combination of genomic and histological techniques, we identified Alphamesonivirus-1 in two horses that succumbed to an acute respiratory syndrome. This represents the first detection of a mesonivirus in vertebrates.

## RESULTS

### Bronchopneumonia and unspecific-viral infection detected in lungs and lymph nodes of two horses that succumbed to an acute respiratory syndrome

The body condition score of the horse carcasses were in the physiological range, with no signs of external trauma observed. The mare was in foal, and the intra-uterus foetus was of normal physiological size and shape. Advanced putrefaction and decomposition of internal digestive organs as late post-mortem changes were apparent. Post-mortem analysis revealed foamy nasal discharge, severe gelatinous subcutaneous oedema of the neck region, and hemorrhagic pleural exudate, while severe pulmonary oedema and thickness of interlobular septa were observed in the lungs of both horses. Splenomegaly, enlargement of bronchial, submandibular, and retropharyngeal lymph nodes (LN), the necrosis of submandibular LN, and petechial hemorrhages of the large intestine mucosa were evident as well (Figure 1A). Histopathology analyses revealed severe alveolar oedema, massive and severe thickness of visceral pleura, areas of bronchopneumonia and hemorrhagic foci in bronchial, submandibular and retropharyngeal lymph nodes (Figure 1B). Both the mare and the foal showed gross and microscopic similar lesions in terms of localization, type, and extension (Figure 1C and 1D).

**Figure 1.**
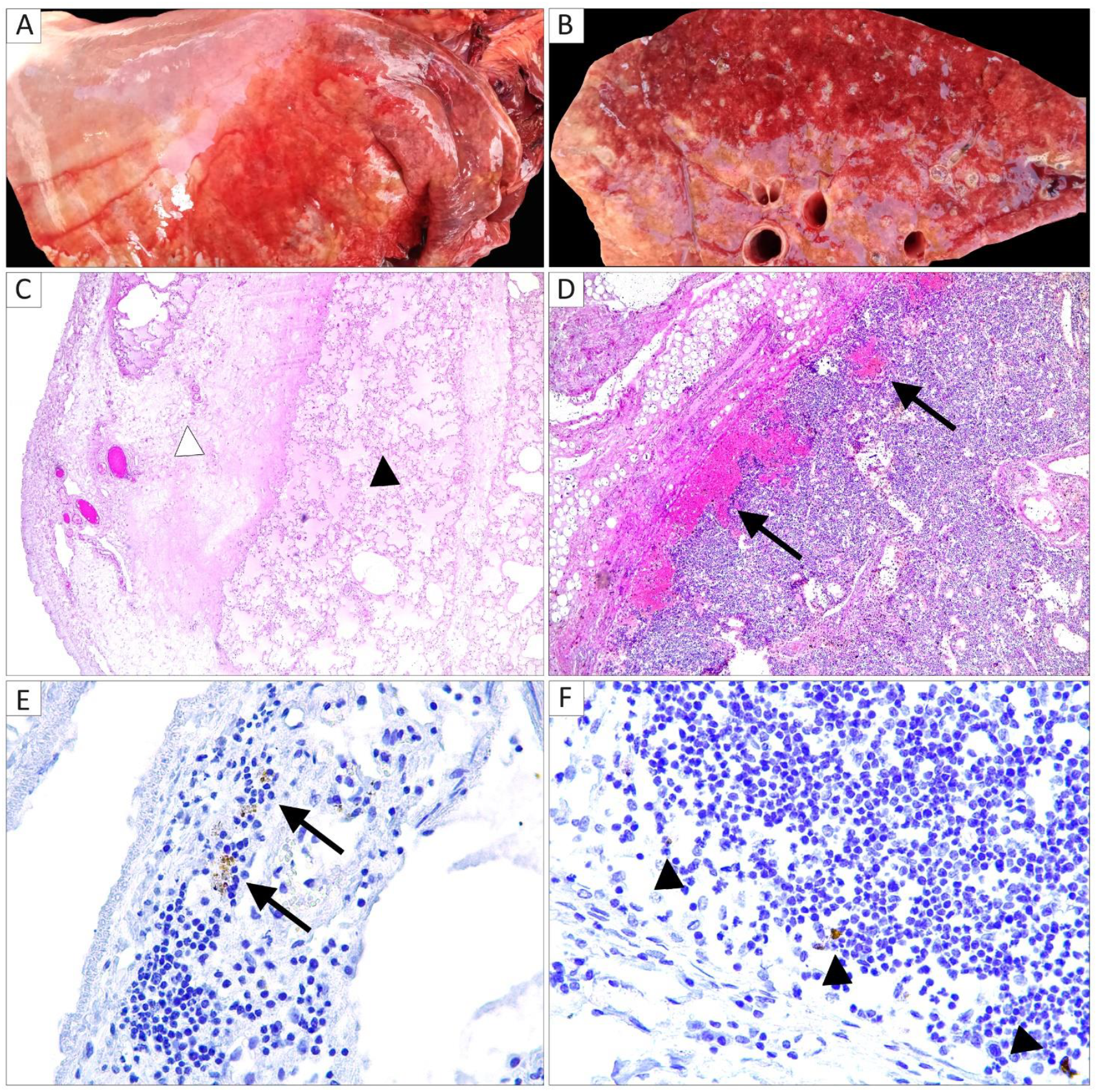
Organ photographs and histological staining of tissues from the horses that succumbed to an acute respiratory syndrome. At gross examination of the lung, the mare showed pleural effusion (A), and on cut section, pulmonary oedema and marked enlargement of interlobular septa (B). Histological analyses (hematoxylin and eosin stain) in the lung demonstrated (C; 50x magnification) thickness of visceral pleura (black asterisk) and diffuse alveolar oedema (white asterisk), and in a bronchial lymph node (D; 100x magnification) sub-cortical multifocal hemorrhages (black arrows). Alphamesonivirus-1 *in situ* hybridization (400x magnification) detected intracytoplasmic signals visual as brown spots in scattered macrophages residing the sub-capsular sinus in a bronchial lymph node (E) and in lung alveoli (F).

All samples tested negative for the common agents responsible for respiratory syndrome in horses: EHV1, EHV4, WNV, USUV, EAV, IAV, AHSV, *Babesia caballi, Theileria equi, Trichinella spiralis* (Table 1). Bacterial growth was not observed when the brain, lung and lymph nodes homogenates were cultured with standard protocols. Anaerobic bacterial growth was observed in spleen, kidney, and liver tissue homogenates likely as result of post-mortem proliferation.

**Table 1.**
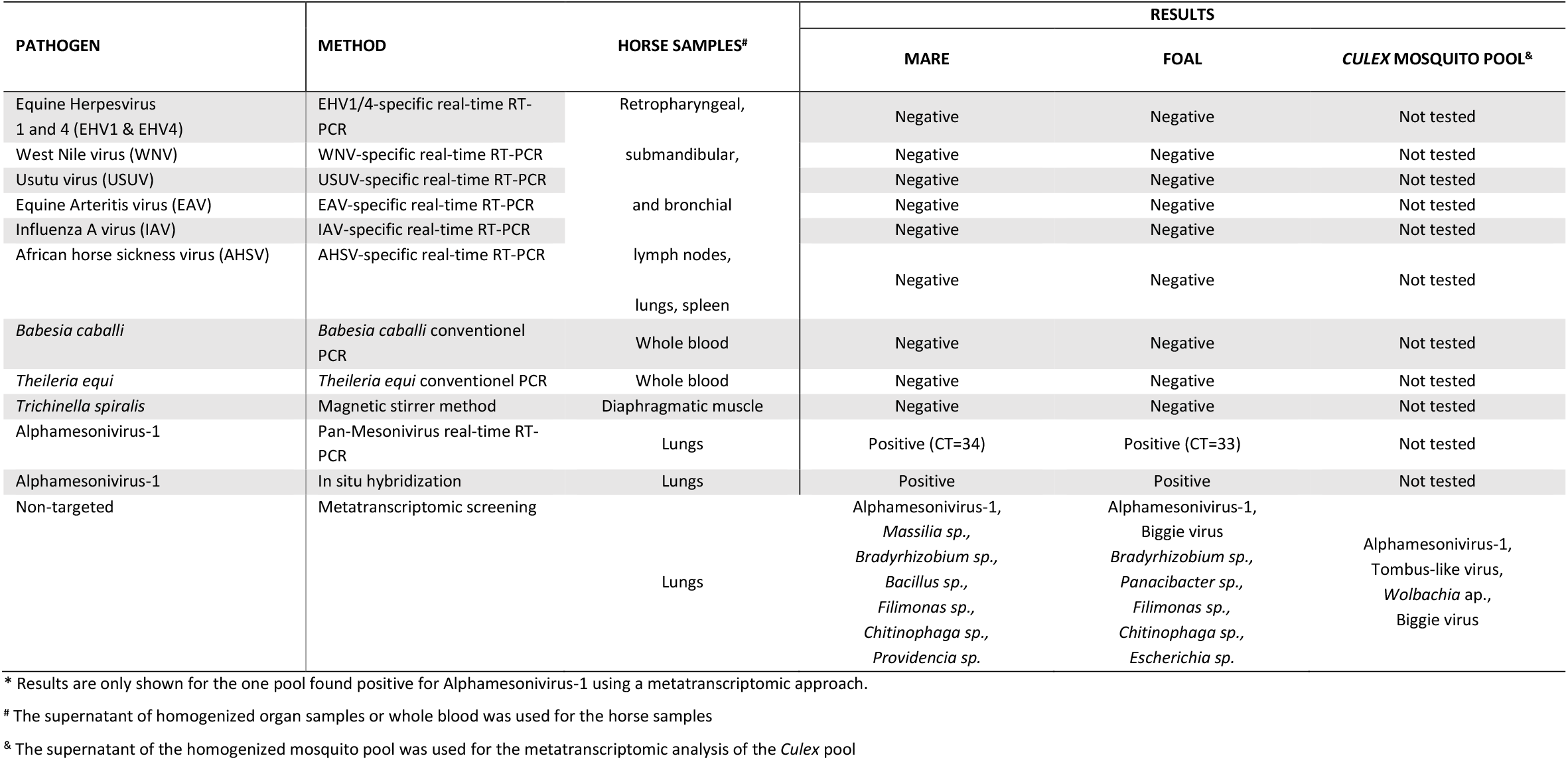
Pathogen detection in horses (mare and foal) and a *Culex* mosquito pool*.

### Identification and confirmation of Alphamesonivirus-1 in horse samples

Metatranscriptomic sequencing produced a total number of 28,231,914 and 26,934,576 raw reads from the foal and mare lung tissue samples, respectively. After quality checks and trimming, the remaining 26,944,930 and 25,958,036 reads were analyzed using the CZID tool: this assigned 14,683 reads (0.73% of all reads, foal) and 746 (0.43%, mare) reads to Alphamesonivirus-1 strain pool 11/2008 (GenBank: MF281710.1) identified from a *C. pipiens* pool in north Italy in 2008. Similar results were obtained by sequential Blast analyses, with RPKM values for Alphamesonivirus-1 ranging from 28.47-54.48 for the foal, and 0.96-1.63 for the mare. Hence, the virus was at considerably higher abundance in the foal than the mare.

A subsequent reference-based assembly of Alphamesonivirus-1 by iVar (1.3.1) produced two consensus sequences with a horizontal coverage (Hcov) of 99% for the foal sample and 57% for the mare sample. Hcov for the mare sample was improved to 87% after re-mapping of reads using the Alphamesonivirus-1 consensus sequence obtained from the foal sample as reference. Mean vertical coverage (Vcov) was 37.22 (min = 1, max = 554) for the foal sample and 5 (min = 0, max= 65) for the mare sample. Pan-mesonivirus real time RT-PCR confirmed the presence of RNA belonging to Alphamesonivirus-1 in both samples and threshold cycles (CT) were 33 and 34 for foal and mare lungs, respectively.

In addition to the equine samples, 10 pools of *Culex* sp. were analyzed with this metatranscriptomic protocol. The taxonomic classification of reads by CZID revealed the presence of reads assigned to Alphamesonivirus-1 species in one pooled sample collected in Sant’Omero municipality (Teramo province, Abruzzo region, a neighboring region of Molise). Deep sequencing of this sample resulted in 30,775,924 raw reads. After quality checks and trimming, the remaining 13,125,482 reads were again analyzed using CZID. As before, the most abundant species was Alphamesonivirus-1 (15.07%, with 1,977,945 reads), followed by arthropod-specific microbial species including *Culex*-associated Tombus-like virus (8.7%), *Wolbachia* bacteria (1.53%), and a member of the *Negevirus* genus of positive-sense RNA viruses (0.23%; see below).

*In situ* hybridization for *Alphamesonivirus-1* in both animals demonstrated intracytoplasmic occurrence of viral RNA in macrophages residing the sub-capsular sinus in a bronchial lymph node and in lung alveoli (Figure 1E and F).

### Presence of additional microbial species

As noted above, besides Alphamesonivirus-1 a variety of other microbial species were identified in the horse samples following metatranscriptomics analysis. Perhaps of most note was the presence of transcripts\ for Biggie virus (*Negevirus*, unclassified positive-sense RNA virus) in the foal (RPKM = 1.63), matching its identification in the *Culex* samples.

In addition, almost 2 million reads from the foal lung sample and 250,000 reads from the mare lung samples were assigned to different bacterial species known to be normally non-pathogenic or environmental contaminants (Table 1). Reads assigned to the *Clostridium* genus represented the highest relative sequence abundance in both the mare (91.09%) and the foal (87.27%). Although the vast majority of clostridial species are non-pathogenic commensal or soil bacteria, *Clostridium perfringens* and *C. botulinum* are responsible for severe diseases of horses^41^. Therefore, mapping analysis was performed using the reference sequence for *C. perfringens* (Genbank: CP009557.1) and for *C. botulinum* (CP063816.1) as indicated by the CZID tool. Mapping analysis for both samples failed, resulting in Hcov of 0.01% (foal) and 0.04% (mare) for *C. perfringens* and Hcov of 0.04% (foal) and 0.03% (mare) for *C. botulinum*.

### Characterization of horse Alphamesonivirus-1

The Alphamesonivirus-1 sequences obtained from the foal and mare samples were identical with the exception of ambiguities due to low coverage (resulting in an overall pairwise nt identity of 98.5%). A total of 21 amino acid substitutions were found between the Alphamesonivirus-1 sequences obtained from the horses and from the *Culex* mosquito pool (Figure 2). Overall, eight non-synonymous mutations were found in ORF1a, and ORF1b each, and six in ORF2a. A phylogenetic tree of Alphamesonivirus-1 was estimated using publicly available complete genome sequences combined with those generated here (Figure 3). This revealed some geographical clustering, with monophyletic groups for virus sequences sampled from South Korea, North America, Australia, and Asia. Notably, the sequences generated in this study formed a well-supported monophyletic group (99% bootstrap support) within a European clade (90% bootstrap support) that includes an Alphamesonivirus-1 sequence identified in Italy in 2008 which falls as the sister-group to the horse and *Culex* pool sequences. Hence, this topological pattern is indicative of the ongoing transmission of Alphamesonivirus-1 in Italy.

**Figure 2.**
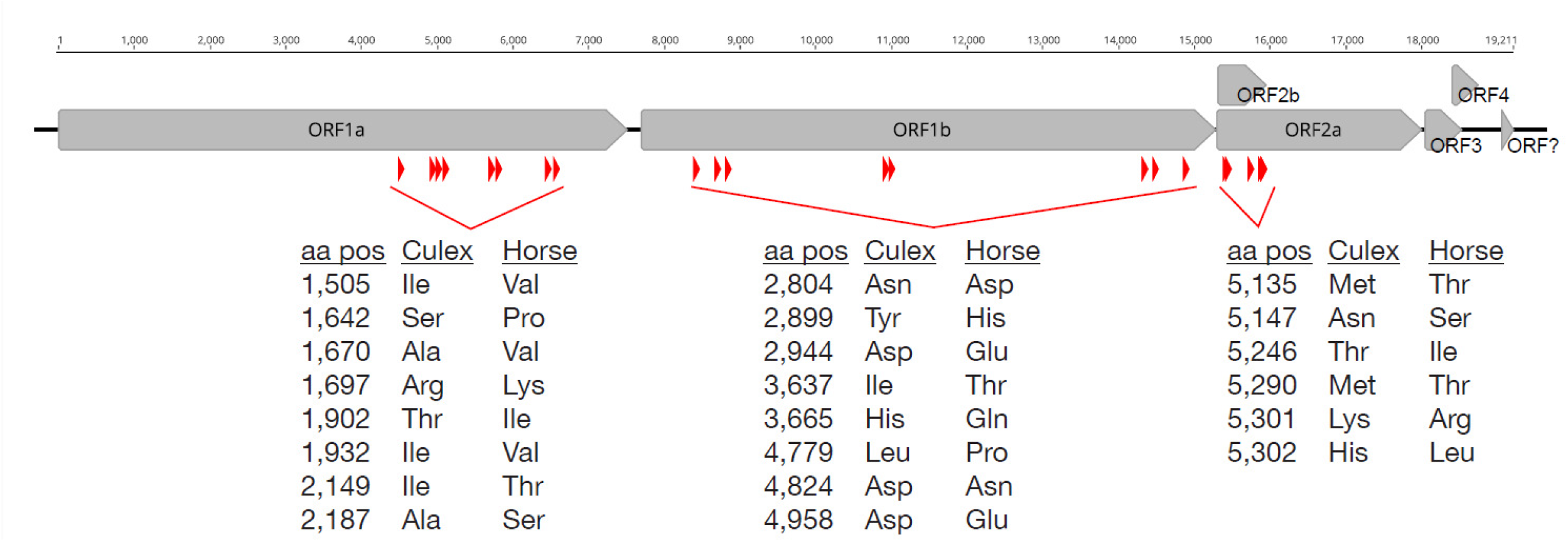
Amino acid substitutions that distinguish the Alphamesonivirus-1 sequences obtained from the horses and from the mosquito pool. A total of 22 amino acid mutations were identified (red arrows) between the Alphmesonivirus-1 sequence from the *Culex* mosquito pool collected in 2022 from Abruzzo region of Italy, and the sequence obtained from the horses with acute respiratory syndrome sampled in 2021. A schematic of the Alphmesonivirus-1 genome is shown.

**Figure 3.**
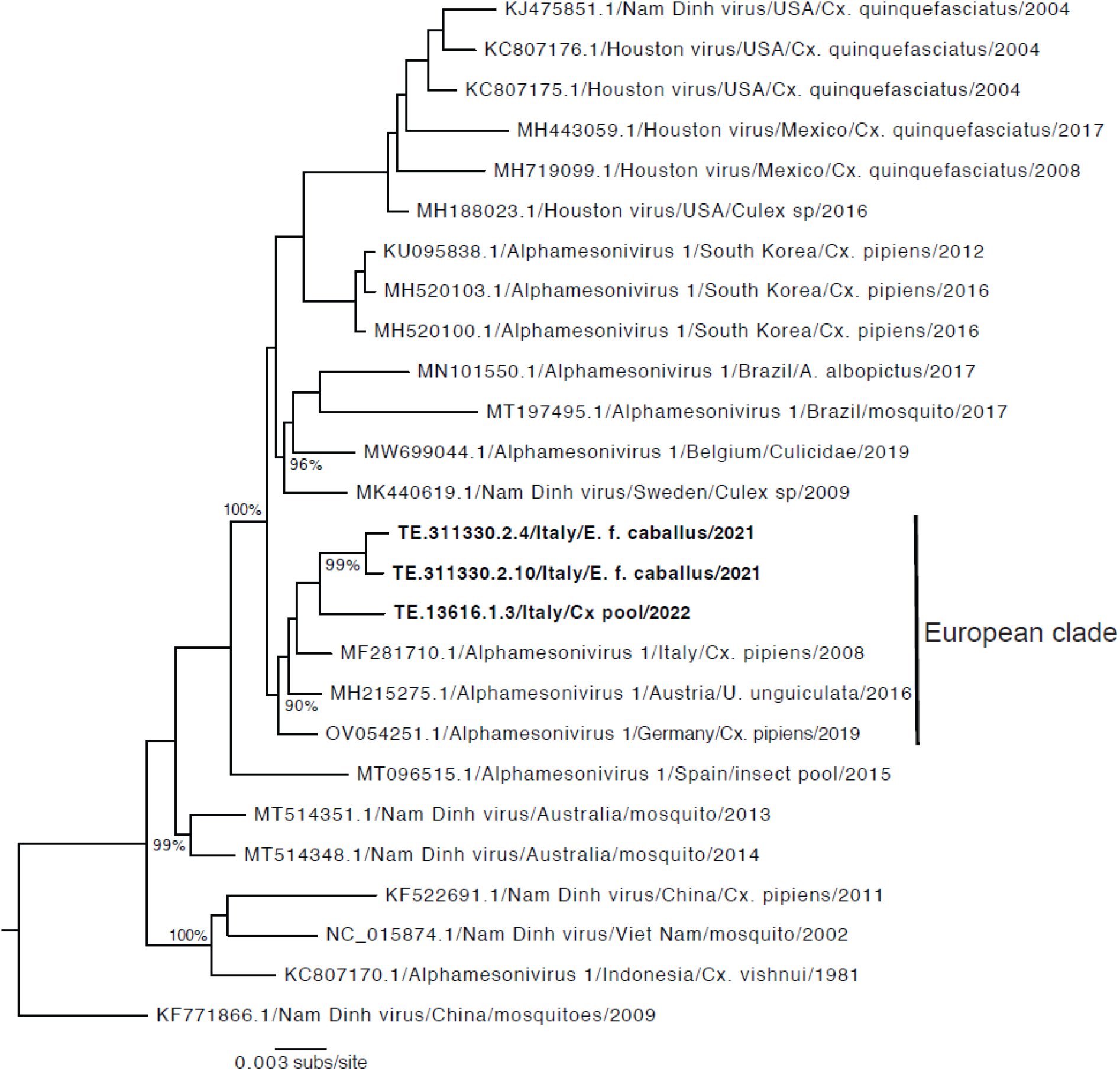
Phylogenetic analysis of the alphamesoniviruses showing the position of the Italian sequences. Maximum likelihood tree of the full genome sequences of alphamesoniviruses with the sequences generated in this study marked in bold (E. f. caballus = horse sequences; Cx pool = mosquito sequences). The European clade of sequences is also marked. The tree is rooted on the first detected Nam Dinh virus sequence (from 2009) and all horizontal branches are drawn to a scale of nucleotide substitutions per site. Bootstrap support values >70% are shown at main internal nodes.

There was no evidence for molecular clock structure in these data (negative correlation coefficient -0.29, R squared 0.08), suggesting that Alphamesonivirus-1 is evolving relatively slowly over the time course of sampling and precluding more detailed molecular clock analysis. This is in marked contrast to other *Nidovirales* such as SARS-CoV-1 and SARS-CoV-2 that have experienced measurable evolutionary rates of ∼10^−3^ nucleotide substitutions per site, per year during their spread through human populations^42^. Although some sporadic *in silico* evidence for limited positive selection across the Alphamesonivirus-1 phylogeny was observed, there was no evidence for any adaptive evolution associated with the Italian viruses (results not shown).

## DISCUSSION

Respiratory infections are important causes of morbidity and mortality in horses. However, the causative agents are frequently unidentified, often misdiagnosed or overlooked. Infections of the upper respiratory tract, both viral and bacterial, are usually diagnosed in weanling and yearling horses, while airway disorders, pleuropneumonia or epistaxis conditions caused by inflammation or exercise are found in horses older than two years. In contrast, recurrent airway disease or neoplasia of the respiratory tract are diagnosed primarily in the middle-aged and older horses^43^. Conditions such as inflammatory airway disease, chronic obstructive pulmonary disease or exercise-induced pulmonary hemorrhage are examples of the pathological conditions affecting respiratory system of horses, whose complex etiology remain uncertain. Nevertheless, viral respiratory infections are the most important causes of respiratory disease in horses worldwide.

Mesoniviruses have no known association with human or animal disease. To the best of our knowledge, this is the first report describing Alphamesonivirus-1 infection in a vertebrate. The presence of this virus in two horses located in the same barn was suggested by metatranscriptomic analysis and confirmed by an Alphamesonivirus-1-specific-PCR and *in situ* hybridisation on infected pulmonary tissues. Phylogenetic analysis revealed that the Alphamesonivirus-1 identified from the mare and the foal from the Molise region of Italy were closely related to a sequence derived from a mosquito pool collected in the neighboring Abruzzo region in 2022. The high genetic similarity between these sequences, and to sequence from a virus identified in north Italy in 2008, suggests that Alphamesonivirus-1 is continuously circulating in Italy.

Microbiological analyses of kidneys, liver, and spleen tested positive for the presence of anaerobic bacteria, yet culture isolation, including the pathogenic *Clostridium perfrigens*, yielded negative results. Observed anaerobic bacterial growth was, therefore, likely present due to post-mortem changes of decomposition and putrefaction of the internal organs.

As no other well-recognized horse pathogens were identified in our analysis, it is tempting to speculate that the Alphamesonivirus-1 infection had a pathogenic role, alone or in association with an undiagnosed pathogen, in the acute respiratory syndromes observed in the horses. However, this will need to be confirmed, and the mechanisms triggering the infection remain unknown. Similarly, it is uncertain whether the virus was maternally transmitted from mare to foal, or whether these animals were simultaneously infected by local *Culex* mosquitoes.

The presence of the Biggie virus (*Negevirus*) in the foal and in the Culex mosquito pool is also intriguing. Biggie virus is associated with *Culex* mosquitoes and was first identified at relatively high abundance in *C. pipiens* and *C. torrentium* from Sweden^44^. Although, in a similar manner to Alphamesonivirus-1, there is no prior evidence for negeviruses in mammalian species, these two viruses were recently found to interact *in vitro* with several co-infecting arboviruses (*Flaviviridae, Togaviridae*, Perib*unyaviridae*) inhibiting USUV and Bunyamwera orthobunyavirus infection^45^. The co-infection of Alphamesonivirus-1 and Biggie virus therefore merits additional attention.

Similarly, we cannot exclude a major contributing role of a cytokine storm following Alphamesonivirus-1 infection for the observed respiratory syndrome, particularly as this is well known in related *Nidovirales* including feline infectious peritonitis virus and SARS-CoV-2^46–49^. COVID-19 pathology was generally characterized as biphasic with an acute phase dominated by active SARS-CoV-2 infection and a post-viral clearance phase dominated by host reparative and immunologic processes^50^. Hence, we cannot exclude macrophage hyperactivation in the two horses since macrophages in the sub-capsular sinus in a bronchial lymph node and in lung alveoli were shown to be positive for Alphamesonivirus-1 by *in situ* hybridization. This scenario is also described in SARS-CoV-2 infected individuals in which CD169+ macrophages were detected in lymph node subcapsular spaces^51^. Macrophages disorders such as secondary hemophagocytic lymphohistiocytosis have been well described in COVID-19 and in other coronavirus infections such as SARS and MERS^52^. Hemophagocytic lymphohistiocytosis is a hyperinflammatory syndrome characterized by a fulminant and fatal hypercytokinaemia with multiorgan failure in humans. In adults, this phenomenon is mostly triggered by viral infections, autoimmune diseases, and neoplasms^53^.

The low viral load found in our study indicated by high ct values and low number of Alphamesonivirus-1 reads calls into the question whether this virus was the causative agent of the fatal respiratory disease in the horses. However, similar circumstances were observed when Schmallenberg virus was first detected in blood samples from infected cows^54^. This virus is transmitted by the female biting midges of the *Culicoides obsoletus complex* and is associated with disease ruminants including fever, fetal malformation, drop in milk production, diarrhoea and stillbirths, becoming a burden for small and large farms^55^.

Despite all attempts with different mammalian and mosquito cell lines (VeroE6 and C6/36), we were unable to isolate the virus. As the experiments undertaken in by Diagne *et al*.^12^ demonstrated the inability of Alphamesonivirus-1 to replicate in the mosquito C6/ 36 cells at 37 °C but at only 28 °C, we performed all isolation attempt (also with mammalian cell lines) at 28 °C. Another limitation of the study is that we were unable to screen a healthy population of horses from the same farm to investigate the presence of Alphamesonivirus-1 RNA genome in the rest of the herd. Furthermore, it would have been beneficial to collect mosquitoes from that farm. Due to the lack of specific serological tools for Alphamesonivirus-1, the local virus circulation and potential seroconversion in horses could not be addressed. Due to the presentation as an acute respiratory syndrome, the gross lesions described, and the advanced state of putrefaction observed in the carcasses, we initially focused only on the lungs and did not perform the Alphamesonivirus-1 PCR and metatranscriptomics on other organ samples.

Overall, the presence of Alphamesonivirus-1 in two horses may provide new insights into the pathogenesis of respiratory diseases of horses, and enhances our understanding of the diversity and evolution of mesoniviruses. The correlation between the pathological condition observed in these horses and the presence of Alphamesonivirus-1 in their lungs represents an important first step in understanding mesonivirus evolution, host range, and its potential to infect and cause disease in other animals than insects. Since the presence and/or replication of Alphamesonivirus-1 in mammalian hosts has not been reported to date, our identification of a supposedly insect-specific virus in a mammalian host clearly necessitates the further *in vivo* investigation and broader surveillance of this virus.

## MATERIALS AND METHODS

### Sample collection and diagnostic approach

In October 2021, an 18-month-old foal and a 7-years-old mare Haflinger horse died unexpectedly due to acute respiratory syndrome in the same farm located in Miranda, province of Isernia, Molise region, Italy. The two carcasses were sent to the *Istituto Zooprofilattico Sperimentale dell’Abruzzo e del Molise* (IZSAM) for necropsy. Lung and bronchial lymph nodes were sampled from each animal, fixed in 10% neutral buffered formalin, routinely processed for histology and stained with Haematoxylin and Eosin (HE). Samples from the retropharyngeal, submandibular and bronchial lymph nodes, lungs and spleens were collected and homogenized in a sterile phosphate-buffered saline (PBS), and then centrifuged. Nucleic acid was extracted from 200 µl of supernatants using the MagMAX CORE Nucleic Acid Purification Kit (Applied Biosystems) on an automatic extractor KingFisher Flex (ThermoFisher Scientific), with an elution volume of 100 μl, following the manufacturer’s instructions. The samples were tested by molecular assays for the presence of RNA/DNA of several respiratory viruses, including EHV1 and EHV4^24^, West Nile virus (WNV)^25^, Usutu virus (USUV)^26^, *Alphaarterivirus equid* (EAV) (VetMax EAV Kit, Applied Biosystems), IAV^27,28^, African horse sickness virus (AHSV)^29^. Whole blood samples were analyzed for the detection of *Babesia caballi* and *Theileria equi* by conventional PCR^30,31^, while diaphragmatic muscle was tested for the presence of *Trichinella spiralis* by means of magnetic stirrer method^32^. All samples were also tested by standard procedures for aerobic and anaerobic bacterial isolations.

### Sample preparation, library construction and metatranscriptomic sequencing

To assist with pathogen identification, RNA purified from lung samples of both horses was processed for metatranscriptomic analysis. After Turbo DNAse (Thermo Fisher Scientific, Waltham, MA, USA) treatment at 37°C for 20 min, total RNA was purified by an RNA Clean & Concentrator™-5 Kit (Zymo Research, Irvine, CA, USA). The RNA obtained was processed using sequence-independent single-primer amplification protocol (SISPA) with some modifications^33^. The amplicons were purified by ExpinTM PCR SV (GeneAll Biotechnology CO., LTD Seoul, Korea), and quantified by Qubit dsDNA HS assay (Thermo Fisher Scientific, Waltham, MA, USA). The samples were diluted to obtain a concentration of 100–500 ng and used for library preparation with the Illumina DNA Prep kit (Illumina Inc., San Diego, CA, USA) according to the manufacturer’s protocol. Deep sequencing was performed on the NextSeq 500 (Illumina Inc., San Diego, CA, USA) using the NextSeq 500/550 Mid Output Reagent Cartridge v2, performing 300 cycles and generating 150 bp paired end reads. Raw sequencing reads underwent quality trimming before adapter removal using Trimmomatic v0.38. Quality control of raw and trimmed reads was performed with FASTQC v0.11.8.

Fastq files were initially analyzed using the Chan Zuckerberg ID (CZID) software (https://czid.org/), an open-source software platform that helps identify pathogens in metatranscriptomic sequencing data after host sequence removal. Following indications on microbial composition provided by CZID, fastq data of both horses were mapped by BWA software package (v.0.7.17)^34^ to Alphamesonivirus-1 reference accession number NC_015668. Alphamesonivirus-1 consensus sequences were obtained by iVar (v1.3.1). Pairedend reads were *de novo* assembled into contigs using MEGAHIT v1.2.9 with default settings. To confirm these initial observations, the assembled contigs were compared to the NCBI non-redundant database (NCBI-nr) using DIAMOND v2.1.6 with an e-value cut-off ≥1E-4^35^. To provide further validation of hits, contigs were screened against the nucleotide database (NCBI-nt) with an e-value cut-off ≥1E-10. Virus abundance was quantified and normalized by contig length using TPM (transcripts per million) and RPKM (reads per kilobase million) metrics as implemented in RSEM v1.3.0.

Within genomic surveillance activities performed at IZSAM in 2022 within the National surveillance system of arboviral diseases, 10 pools of *Culex* sp. were collected in different parts of Teramo province, Italy (in the Abruzzo region, a neighboring region to Molise). These samples also underwent metatranscriptomic analysis using the protocol described above.

### Specific Pan-Mesonivirus real time RT-PCR

The presence of Alphamesonivirus-1 was confirmed by real-time RT-PCR assay using primers and probes as described in Diagne *et al*. (2020). PCR reactions were prepared with 10x GoTaq Probe qPCRMaster Mix (Promega) containing final concentrations of 0.5 mM forward primer, 0.5 mM reverse primer, 0.25 mM TaqMan probe, 5 µl of double-strand cDNA obtained after SISPA protocol, and nuclease-free water up to 20 µl reaction volume. Real-time RT-PCR reactions were performed on a QuantStudio 7 Flex Real-Time PCR System (Applied Biosystems) in fast mode with the following settings: initial denaturation at 95°C for 20 sec, followed by 40 cycles of denaturation at 95°C for 1 sec and annealing/extension at 55°C for 20 sec.

### RNA *in situ* hybridization

Lung and bronchial lymph nodes derived from the foal and mare were subjected to an RNA *in situ* hybridization performed by RNA scope analysis platform (RNAscope 2.5 HD Assay – BROWN kit, Biotechne) following the manufacturer’s instructions. To detect viral RNA, sections were incubated with an *ad hoc* probe designed and manufactured commercially (ACDbio, Bio-Techne, USA). The probe was designed to detect Alphamesonivirus ORF1a (GenBank: MT096515.1) targeting nt 1084-2098. The endogenous housekeeping gene Ubiquitin C (UBC) was used as positive control to assess both tissue RNA integrity and assay procedure. Slides were counter-stained with Mayer’s Hematoxylin (Bio-Optica, Italy) and mounted with Eukitt® mounting media (Bio-Optica, Italy) before analyzing on a Zeiss Axio Scope.A1 microscope (Carl Zeiss Microscopy GmbH, Göttingen, Germany).

### Phylogenetic and evolutionary analysis

Nucleotide sequences representing the full-genome of Alphamesonivirus-1 were obtained from NCBI (accessed 5 June 2024; n=52) and combined with two reference sequences (NC_015874.1 and NC_015668.1) and the two sequences isolated from both horses and from the mosquito pool. A multiple sequence alignment was performed in Mafft v7.450 using the FFT-NS-i x1000 algorithm (Katoh and Standley, 2013). The alignment was manually inspected in Geneious Prime 2021.1.1 (https://www.geneious.com) for accuracy. Sequences with extensive genetic diversity were removed. A maximum likelihood (ML) phylogenetic tree was then estimated for the final data set of 43 full-genome sequences using RAxML v 8.2.11 (Stamatakis, 2014) implementing a gamma time reversible +Γ model of among suite rate heterogeneity (GTR+Γ) nucleotide substitution model and 200 bootstrap replicates.

To assess the extent of temporal (i.e., clock-like) structure in the data we performed a regression of root-to-tip genetic distance on the ML tree against date (year) of sampling using the TempEST method (Rambaut et al., 2016). The lack of temporal structure (see Results section) precluded additional analyses of evolutionary dynamics.

To test for the presence of positive selection (i.e., adaptive evolution), especially on the amino acid substitutions associated with the viruses in the horses, we utilized the FUBAR^36^, MEME^37^, BUSTED^38^ methods in HyPhy/Datamonkey^39,40^ that explore various distributions of the numbers of nonsynonymous (d_N_) and synonymous (d_S_) substitutions per site.

## Data availability

The two Alphamesonivirus-1 consensus sequences obtained in this study from the Haflinger mare and foal, as well as the sequence from the *Culex* mosquito pool, were deposited in the NCBI/GenBank database under accession numbers PP961236, PP961235, and PP961237, respectively. Metatranscritptomic data are available on the Sequence Read Archive (SRA) under bioproject number PRJNA1126112.

## ACKNOWLEDGEMENTS

We acknowledge Dr. Addolorato Ruberto (Istituto Zooprofilattico Sperimentale dell’Abruzzoe del Molise) for horse necropsy.

## Authorship contributions

**Lucija Jurisic**: Conceptualization, Data curation, Writing – original draft

**Heidi Auerswald**: Writing - review & editing

**Maurilia Marcacci**: Conceptualization, Investigation, Writing – original draft

**Francesca Di Giallonardo**: Formal analysis, Visualization

**Laureen M. Coetzee**: Investigation, Methodology, Formal analysis

**Valentina Curini**: Formal analysis

**Daniela Averaimo:** Formal analysis

**Giovanni Di Teodoro**: Methodology, Software, Validation, Formal analysis, Investigation, **Ayda Susana Ortiz-Baez**: Methodology, Software, Validation, Formal analysis, Data curation **Cesare Cammà:** Writing – review & editing

**Juergen A. Richt**: Conceptualization, Writing – review & editing

**Edward C. Holmes**: Conceptualization, Writing – original draft, Writing – review & editing **Alessio Lorusso**: Conceptualization, Writing – original draft, Writing – review & editing, Funding acquisition, Supervision, Project administration

## Declaration of competing interest

The authors declare that they have no known competing financial interests or personal relationships that could have appeared to influence the work reported in this paper.

### Preprint to bioRxiv

All authors agree on posting this article to bioRxiv

